# Exploring the Educational Potential of Instagram in the New Generations: A Study on the Impact of “Adopt a Bacterium”

**DOI:** 10.1101/2024.10.01.616198

**Authors:** Bárbara Rodrigues Cintra Armellini, Lara Nardi Baroni, Samantha Carvalho Maia Brito, Ana Carolina Ramos Moreno, Robson Francisco de Souza, Camilo Lellis-Santos, Rita de Cássia Café Ferreira

## Abstract

In 2013, the “Adopt a Bacterium” project was created, which originally proposed the use of the social network Facebook® as an educational tool to engage students in learning microbiology through personalized posts. However, the generational alignment dynamic presents contemporary challenges for educators. Given the relevance and success of the project in recent years, the use of an alternative social network is sought for the application of the methodology. In the present study, we describe the use of the social network Instagram as a tool to understand the impacts of the “Adopt a Bacterium” project on microbiology learning among new generations of students. To verify the effectiveness of the methodology, the following steps were taken: i) voluntary and anonymous questionnaires with an open-ended question about knowledge of the adopted bacterium were applied before the start of the project (Q1), at the end of the project (Q2), and five months after the end (Q3), to Biomedical Sciences students in the years 2020, 2021, and 2022; and ii) a project evaluation questionnaire was administered to students. The data were analyzed using thematic categorization, calculation of the richness of student’s discourse through the Shannon Diversity Index, and the creation of word clouds to visualize student suggestions, as well as the use of Python language tool to demonstrate citations of specific themes, comparing Facebook and Instagram as Virtual Learning Enviroments. The results revealed an improvement in student learning, as demonstrated by the increase in the frequency of content categories related to bacteriology and the richness of discourse, indicated by higher Shannon Diversity Index values in Q2 and Q3 compared to Q1. In conclusion, the results demonstrate that the methodology of the “Adopt a Bacterium” can be applied in different virtual environments, such as the social network Instagram® resulting in increased learning and student interest in Microbiology.

## Introduction

The traditional teaching model, that is, the one based on lecture-based teaching classes, has been under discussion for decades, with indications that its reformulation is necessary. This is mainly because this educational model is centered on the figure of the teacher, considered the holder and transmitter of knowledge to students, who are seen as passive recipients. The autonomy of the student is advocated by many authors, who argued that students should be actively present in the classroom, developing autonomy to learn and thus constructing their own knowledge^1^. This idea about the role of the student is strongly supported by the works of Jean Piaget^2^ and David Ausubel^3^ who claimed that by taking on a central role and correlating the concept to be learned with their prior conceptions, students would effectively and enduringly build their knowledge.

These discussions, with these and other educational theorists, form the basis for the development of what are now known as active learning methodologies, which involve methodologies that envision this central role of the student in the classroom. There are numerous active learning methodologies that can be developed both in high school and higher education, and a commonality among most of them is the use of the Internet for at least one of the planned activities, as it is extremely present in the lives of everyone, including students and teachers. For this reason, considering the use of Digital Information and Communication Technologies (DICTs) for educational purposes is interesting, precisely because it brings students closer to the content to be studied. The Internet is becoming increasingly prominent and important in the lives especially in Brazil which, with India and China, accounts for about 70% of global access, ranking 4th in this regard^4^. Furthermore, approximately 70% of the Brazilian population is connected to the Internet in some way^5^.

There are many ways to use the Internet for educational purposes, and one of these possibilities is the use of social networks as Virtual Learning Environments (VLEs), which is quite interesting considering that they are widely used, especially in Brazil, where, according to the site We Are Social, about 60% of the population is connected to them. Furthermore, several studies have shown that their use contributes to student learning, both in higher education and high school^6,7^. There are numerous possibilities for incorporating the online environment into the educational context, one of which is Blended Learning, which has been studied for some time and presents itself as an innovative methodology^8^ precisely because it consists of a mix between traditional classes and the virtual environment, aiming to make learning continuous and occur in multiple environments^9^. With it, the goal is to add value to the learning environment by incorporating resources from the virtual environment, like hyperconnectivity, immediatism and contemporary belonging^10^, although traditional teaching remains the cornerstone of student education^11^.

Given the relevance and importance of the Internet and social networks to the population, the #Adopt Project was created in 2013, initially using Facebook® as an educational tool for teaching Microbiology. Facebook® was selected at the start of the project because it was one of the most widely used social networks by the population, including teachers and students^12^, currently used by 14.5% of the world’s population, according to the site We Are Social. It is also interesting because it promotes discussions and enhances the teacher-student relationship^13^, enabling greater interactions between them^14^. However, the migration to Instagram® was required because its growing importance, now accessed by 14.4% of the world’s population, according to the We Are Social platform. Additionally, it is on this social network that the young audience can be found, as about 65% of its users are less than 35 years old.

Initially, “Adopt a Bacterium” was developed for higher education with its “Adopt a Bacterium” version^15^ and later for high school with its “Adopt a Microorganism” version, whose use during the emergency remote teaching period caused by the COVID-19 pandemic was effective and relevant for the teaching-learning process of participating students^16^.

“Adopt a Bacterium” and its activities provide students with the opportunity to develop their autonomy and important competencies, such as “Learning to Learn,” one of the four pillars of education for the 21st century^17^. With this, students learn how and where to seek relevant information that helps them learn various concepts and content. Furthermore, discussions on social networks and the deepening of topics are led by postgraduate and undergraduate students who act as mediators. This role is very important as it allows the development of important skills for these mediators, enhancing their professional development^18^.

The pandemic period was very impactful for various sectors of society, especially the educational sector, which faced significant concerns about student engagement and interest, as they were forced to attend classes and carry out academic activities remotely^19,20^. Additionally, some studies show that there was a loss of learning during this period, making the “Adopt a Bacterium” an interesting methodology to engage students during this complex time. Verifying whether there was learning during this period and its different phases (emergency remote, remote, and return to in-person model) is also important to understand the impact of this time, as well as the methodology, on students.

Thus, the objective of this work is to verify how the “Adopt a Bacterium” impacts the learning among higher education with the proposal to expand the “Adopt a Bacterium” to multiple social media platforms, particularly with Instagram®, and its effectiveness both to online (during the pandemic period) and presential formats.

## Material and Methods

The methodology of the present project will be described according to: the development of “Adopt a Bacterium” in the years 2020, 2021, and 2022, with the first two being developed in a remote model and the last in an in-person teaching model; data collection; analysis of the frequency of category citations in the questionnaires; creation of word clouds and calculation of the Shannon Diversity Index^21^. The recruitment for the study started on August 5, 2020, and was completed on April 20, 2023 and the participants signed a written consent form.

### The Development of “Adopt a Bacterium”

“Adopt a Bacterium” was developed with the Biomedical Sciences and Fundamental Health Sciences classes in the Bacteriology course during the years 2020, 2021, and 2022 that were chosen because they were the years of the COVID-19 pandemic (2020 and 2021) and post-pandemic (2022). In this project, students were divided into groups and adopted a bacterial species or genus. In 2020, the adopted bacteria were *Corynebacterium diphtheriae, Mycobacterium tuberculosis, Neisseria sp*., *Streptococcus sp*., *Treponema pallidum*, and *Yersinia pestis*. In 2021, they were *Escherichia coli, Mycobacterium tuberculosis, Streptococcus sp*., *Treponema pallidum*, and *Vibrio cholerae*. Finally, in 2022, the adopted bacteria were *Mycobacterium tuberculosis, Escherichia coli, Staphylococcus sp*., *Streptococcus sp*., and *Pseudomonas sp*. The bacteria are selected year by year according to the direction of the discipline, but despite the differences, the levels of difficulty and demands have been homogeneous over the years.

During the class period, students made posts on social networks (Facebook® in 2020 and Instagram® in the following years) relating the topics covered in the theoretical and practical classes of the course to the bacteria they adopted. The posts were only focused on the topics that needed to be addressed, but not on how this approach should be done. These posts were corrected, discussed, and further explored with the mediators of each group. At the end of this period, the students prepared a seminar for their peers to share the knowledge acquired throughout the posting period. The aim was for them to discuss the main characteristics of their bacteria creatively and interactively, thus spreading this knowledge among their peers.

Additionally, promotional materials about the adopted bacteria were produced, with the main objective of sharing the knowledge acquired throughout the project with the general public. Finally, a group exam was conducted to evaluate the students in a more playful manner, in line with the goals of the #Adopt Project. The questions, developed by the mediators included clinical cases and word searches, for example.

### Data Collection

For data collection, three voluntary and anonymous questionnaires were administered to examine the concepts that students learned throughout the project. Each questionnaire contained only one open-ended question: “What do you know about the adopted bacterium?” The first questionnaire (Q1) was given before the start of the project to assess the students’ prior conceptions about the adopted bacterium. The second questionnaire (Q2) was administered at the end of the project, before the group exam, to gather information on the knowledge students gained during the #Adopt project. Finally, the third questionnaire (Q3) was given five months after the end of “Adopt a Bacterium” to assess the retention of knowledge acquired during the project.

### Analysis of Category Frequency in Questionnaires Q1, Q2, and Q3

The students’ responses were analyzed according to the themes they addressed and were categorized^22^ into the following categories: none, morphology, genetics, pathogenicity, treatment, prevention, metabolism, ecology, taxonomy, reproduction, life cycle, examples, social impact, and others. Conceptual errors made by students were also assigned to these categories. At this stage, only the presence or absence of each category in each response was considered, without counting the frequency of category appearances within a single response. Subsequently, this individual frequency was counted to determine how often each theme was addressed in each response given by the students in the questionnaires. These data were plotted in Excel® tables, and for the overall frequency, a graph representing the percentage frequency of each theme in each questionnaire (Q1, Q2, and Q3) was generated.

To concisely represent variations in the use of terms related to the themes Morphology, Pathogenicity, and Metabolism, the frequencies of words used by students associated with each theme were collected throughout the year in three different questionnaires and processed in the IPython environment using the Pandas library. The frequency table was converted into tables representing interactions between questionnaires, quantified by the difference in the absolute frequency of words in each one. The interaction tables were loaded into the Cytoscape environment and formatted as networks where each node represents all responses in a given questionnaire and year provided by a single group of students (bacterial species). If the number of words for a given theme decreased in a subsequent questionnaire in the same year, a dotted line was used to represent this variation. In each graph, the size of the node means the number of times words related to the theme was mentioned.

### Calculation of the Shannon Diversity Index

The Shannon Diversity Index^21^ originates from the field of Ecology, where it is used to understand the dynamics and diversity of a community^23,24^. It is also used in Microbiology to understand microbiome dynamics^25,26^. Its use in the field of education has proven feasible, as it allows for systematic analyses and inferences about the knowledge acquisition process and student performance^16^.

The individual frequencies of citations for each thematic category were used as the basis for calculating the Shannon Diversity Index^21^, following the formula:

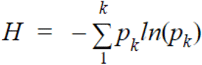

Where p_k_ is the frequency of appearance of each topic in the responses of each group; and k is the total number of different contents that appeared in the responses.

For the overall data, the sum of the individual frequencies of all groups was used, while for the data separated by group, the sum of the individual frequencies of each group separately was used.

### Ethics

This project is approved by the Research Ethics Committee of Human Beings (CEPSH ICB-USP) with CAAE nº 51764021.0.0000.5467.

## Results

In 2020, there were 36 responses in Q1, twenty-two in Q2, and twenty in Q3. However, one response to the initial questionnaire was disregarded for analysis per group, as the student did not identify the bacterium adopted by their group, and their response did not provide any information about it (“I know the morphology” is an example of the response given), rendering its identification and consequent analysis impossible. In 2021, there were forty-four responses in Q1, twenty-eight in Q2 and thirteen in Q3. Finally, in 2022, there were thirty-six responses in Q1, thirty-nine in Q2 and eighteen in Q3. In this year, two responses from the *Mycobacterium tuberculosis* retention questionnaire were disregarded because, similarly to the above-described case, they provided very vague responses that prevented their analysis and categorization.

Initially, a general analysis was conducted of the contents addressed by the students in their responses, categorized into thematic categories based on the core meanings^22^, to understand the general profile of Microbiology contents mentioned by them in their responses, as well as the change in this citation pattern throughout the #Adopt project.

For 2020, it was initially observed that students’ responses in Q1 were generally shorter and more concise, with half of them (50%) related to the pathogenicity of the bacteria adopted by the groups. Additionally, a good portion of them (36.1%) claimed to have no prior knowledge about them (Figure 1A). In 2021, on the other hand, the majority of students (65.9%) claimed to know information about the pathogenicity of the adopted bacteria, followed by approximately 47.72% of them who claimed to know their morphology. In this year, only 15.9% claimed to have no knowledge about the bacterium adopted by their group (Figure 1B). Finally, in 2022, a large portion (47.22%) declared having basic knowledge about the pathogenicity of the adopted bacterium, followed by students (30.55%) who declared having no knowledge about it (Figure 1C).

**Fig 1.**
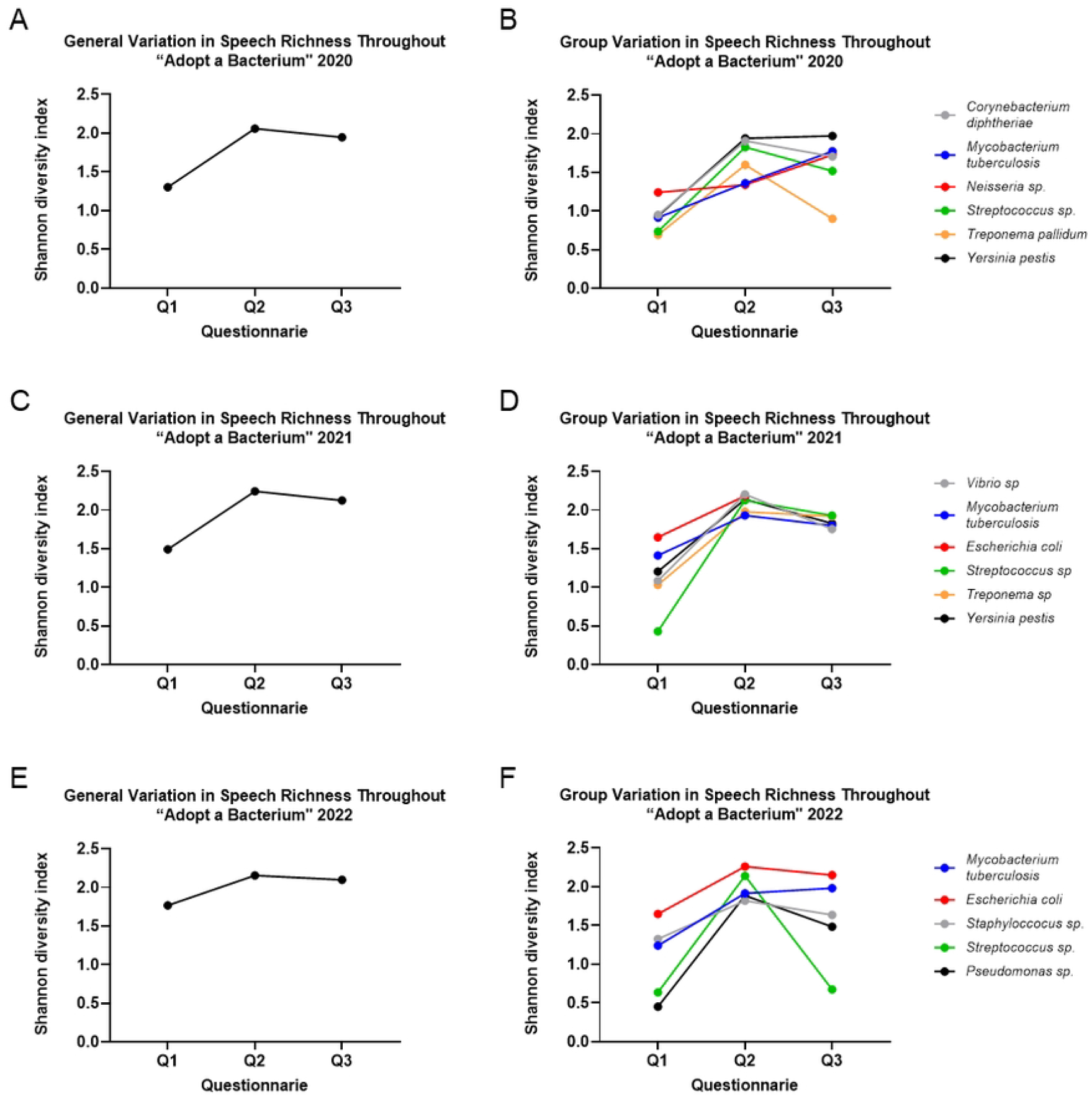
Graphs showing the most addressed contents by students of Biomedical Sciences and Fundamental Health Sciences in their responses to the applied questionnaires, with **[A]** representing the questionnaires answered in 2020 (Q1=36; Q2=20; Q3=22); **[B]** the questionnaires answered in 2021 (Q1=44; Q2=28; Q3=13); and **[C]** the questionnaires answered in 2022 (Q1=36; Q2=39; Q3=18). Q1 is represented by gray color, Q2 by orange, and Q3 by blue.

Regarding the second questionnaire (Q2), in all years, the responses were much longer and elaborate, covering various aspects of the adopted bacteria. In 2020, the majority of students (95.45%) mentioned morphology or aspects related to this theme in their responses, followed by themes of bacterial pathogenicity and metabolism (81.81% in both) (Figure 1A). In 2021, the most focused on aspects related to bacterial morphology and pathogenicity (Figure 1B). In 2022, on the other hand, all (100%) mentioned bacterial morphology in their responses, while a large portion of them (94.87%) also mentioned aspects of its pathogenicity (Figure 1C).

Finally, in the retention questionnaire (Q3), it was observed that the responses were shorter and more direct compared to the previous one, but still more elaborate than those observed in the questionnaire about prior conceptions. In this questionnaire, it was observed that, in 2020, the majority of students (90%) mentioned morphological aspects of the adopted bacteria, followed by citations about bacterial pathogenicity (85%) (Figure 1A). In 2021, on the other hand, all (100%) mentioned bacterial morphology, while 92.3% mentioned aspects related to its pathogenicity (Figure 1B). Finally, in 2022, the same pattern was observed, with all students (100%) focusing on morphological aspects of the adopted bacterium, closely followed (88.88%) by citations related to its pathogenicity (Figure 1C).

This categorization was also performed individually, reflecting the frequency with which each thematic was addressed by each student in their responses, allowing for the calculation of the Shannon Diversity Index^21^.

Regarding the themes, variation in the usage of terms related to the most cited ones in the respective years was observed: morphology, metabolism, and pathogenicity, as demonstrated in Figure 2.

**Fig 2.**
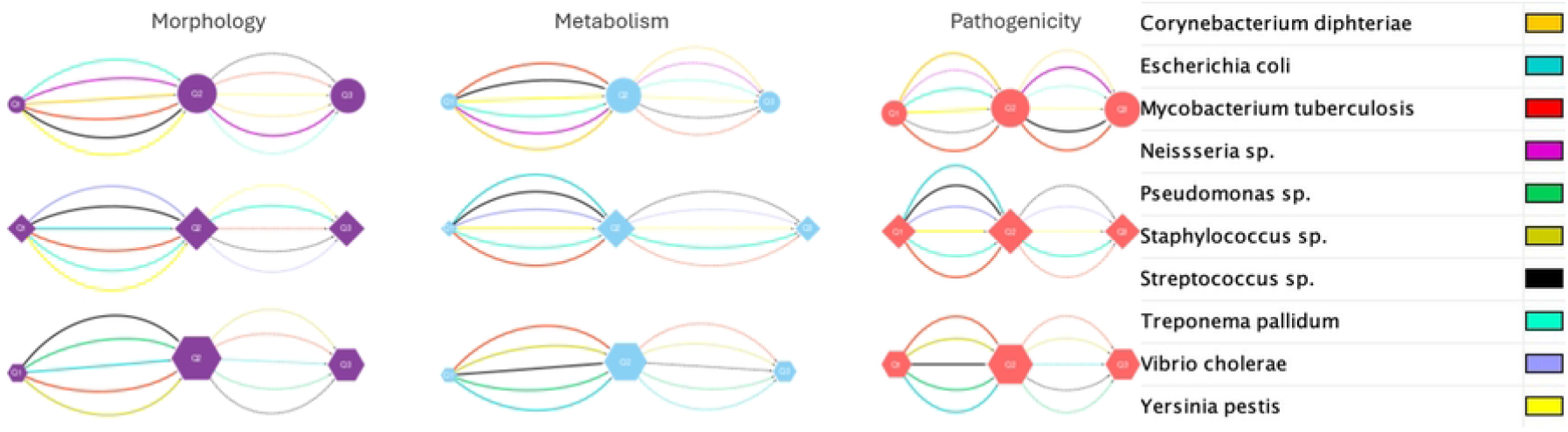
Graphs showing the variation in the usage of terms related to the themes of morphology (purple), metabolism (blue), and pathogenicity (red) by students in their responses to the three applied questionnaires: Q1 (leftmost node in each graph); Q2 (middle node in the graphs); and Q3 (rightmost node in each graph). In each graph, the size of the node means the number of times words related to the theme were mentioned.

It can be observed that all the themes analyzed here undergo an increase in their citation frequency by students between Q1 and Q2, the questionnaire with the largest node size. Between Q2 and Q3, there is generally a decrease in citations, leading to a reduction in node size, especially for the theme of metabolism. However, there is still a high citation frequency when compared to questionnaire Q1.

The richness of students’ discourse in their questionnaire responses was calculated using the Shannon Diversity Index^21^, aiming to assess how it varies throughout the stages of #Adopt and in each group, as from it, inferences can be made about their learning^16^.

For the 2020 class, there is a general increase in discourse richness in Q2 compared to Q1, with a slightly reduced richness in Q3, although it remains higher than that of Q1 (Figure 3A). This general trend repeats for some groups, but those of *Mycobacterium tuberculosis* and *Neisseria sp*. showed higher richness in Q3 than that observed in Q2, as seen in Figure 3B. Additionally, the *Treponema pallidum* group showed a more pronounced decrease in Q3 compared to Q2, although it still remains higher than the richness observed in Q1 (Figure 3B).

**Fig 3.**
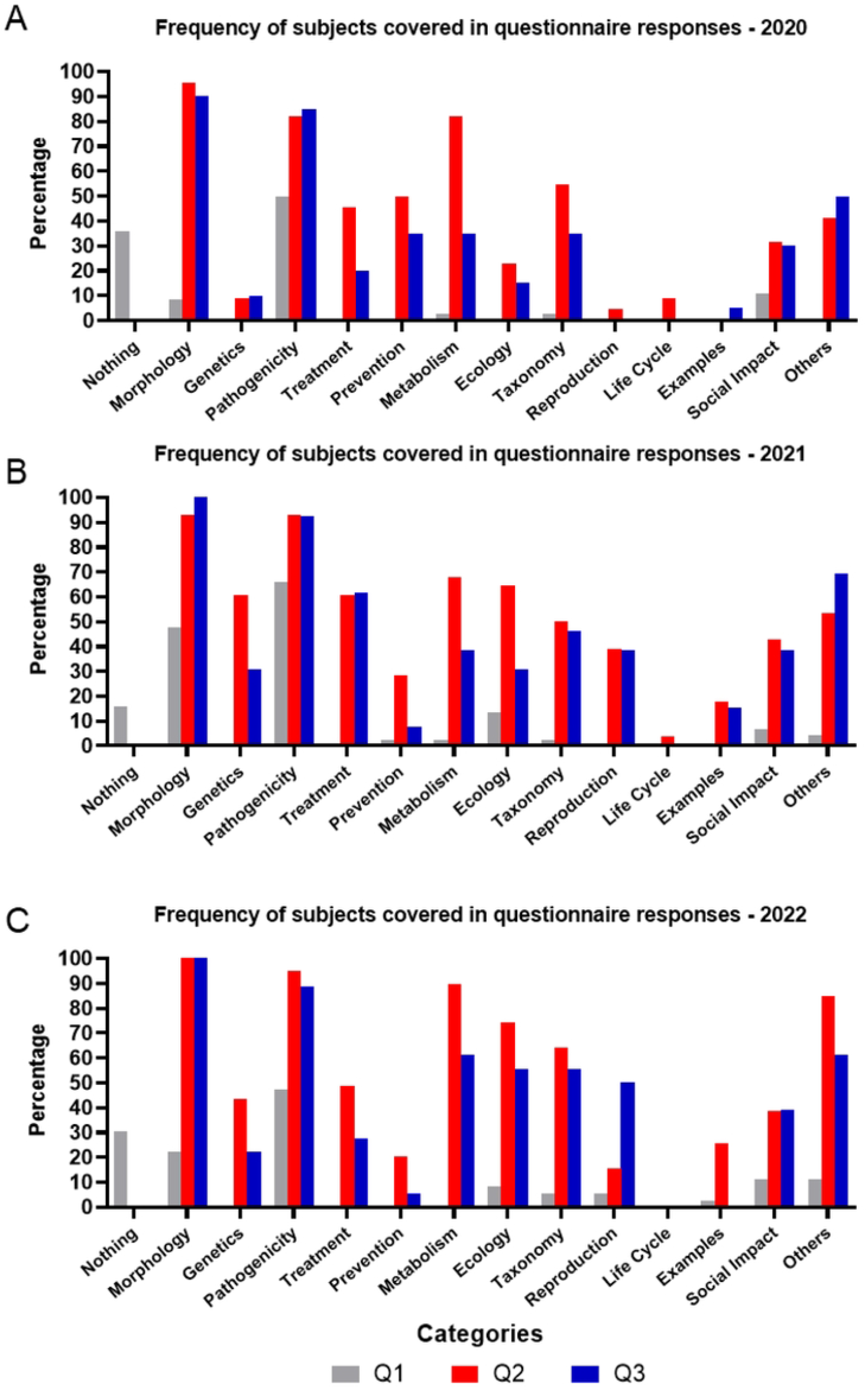
Charts demonstrating the variation of the Shannon Diversity Index throughout the questionnaires applied during the “Adopt a Bacterium” project. The higher the index value, the greater the richness of the students’ responses. **[A]** Index calculated generally for the responses given in 2020; **[B]** Index calculated per group for the responses given in 2020; **[C]** Index calculated generally for the responses given in 2021; **[D]** Index calculated per group for the responses given in 2021; **[E]** Index calculated generally for the responses given in 2022; **[F]** Index calculated per group for the responses given in 2022.

In 2021, overall richness follows a pattern of variation very similar to that described for 2020 (Figure 3C). The same applies to the groups, most of which repeat this observed pattern, except for the *Streptococcus sp*. group, which showed an increase in richness observed at the end of the project (Q2) (Figure 3D), unlike the other groups this year.

Finally, in 2022, the richness varies generally in a manner similar to that described for the two previous years, although the decrease in richness in Q3, compared to Q2, is much less pronounced than in the previous two years (Figure 3E). This pattern repeats for the groups, although the decline in Q3 is more pronounced in most of them (Figure 3F). The major exceptions are the groups of *Mycobacterium tuberculosis* and *Streptococcus sp*., as the former shows an increase in richness compared to Q2, and the latter, in turn, shows an extremely steep decline in richness in Q3, but still higher than that of Q1, as demonstrated in Figure 3F.

## Discussion

These work shows that the “Adopt a Bacterium” is an effective methodology for teaching Microbiology in online and presential formats. The methodology positively impacted students’ learning, increasing the richness of their speech, increasing the themes mentioned and vocabulary on the topic. Besides that, it can be expanded to new platforms, respecting the particularities of each one, according to data obtained from the transition between Facebook and Instagram, present in this work. Its efficiency as a teaching methodology has been demonstrated by learning indicators obtained both through responses to the applied questionnaires and by student grades on proposed activities and informal conversations in the classroom.

The change in social network was necessary due to the change in digital behavior of students, who, as they are part of a new generation, are expanding the range of virtual environments in which they are present, especially because people tend to use social networks in which their friends and closest contact network are present and active^27^. Besides that, they are more motivated and engaged when they are already familiar with the VLE they are using^28^. In this context, understanding whether this change would imply significant losses in learning is necessary and important for the progress and effectiveness of “Adopt a Bacterium”.

The richness of students’ discourse in 2020, when Facebook was used as a VLE, is quite similar to the pattern observed in 2021, when the platform is Instagram. In addition, the variation in the use of terms shown in Figure 2 is also similar between the three years. This similarity in the data patterns allows us to infer that the change in platform and, consequently, the change in VLE does not appear to have affected student learning. Besides that, the similar pattern of the data allows us to infer that there has been consolidation of learning in a very similar way, as the values of the Shannon Diversity Index^21^ are similar in all three years.

An increase in richness is observed throughout the project, with the values of Q2 surpassing those of Q1, indicating that concepts were learned during #Adopt. In the case of Q3, a decrease in its values is expected when compared to Q2, as it is administered five months after the end of the project and students have not, a priori, had contact with this content again. Nevertheless, a value higher than that observed in Q1 is noted, indicating the retention of learned concepts and allowing us to infer that there was indeed consolidation of learning for these students in all scenarios, including that of the pandemic. This information is corroborated by the data observed in Figure 2 for the three years analyzed.

The more pronounced drops in the richness of discourse between Q2 and Q3 are mainly due to the decrease in the number of responses to the questionnaire, especially in the *Treponema pallidum* group in 2020 (Figure 3B) and *Streptococcus sp*. group in 2022 (Figure 3F). This is an expected difficulty since this questionnaire is administered five months after the end of the project, when most students are already taking other courses, often in other departments, which makes it difficult to reach them.

Facebook has a greater focus on text, so the posts were longer and more textually elaborate. On the other side, Instagram is more focused on images, allowing for the exploration of a creative side related to images, drawings, videos, etc^29^. This focus is quite evident when comparing the posts from these years, but the data suggests that there was no impact on concept learning only a change in their presentation format, which is important since young people are increasingly migrating between social medias quickly, and for the methodology to remain attractive and motivating for them, it is necessary to keep up with them in this regard.

The frequency of citations of the themes by students on Facebook (2020) is quite similar to that observed on Instagram, especially in questionnaire Q2, which is a strong indication that, again, the change in VLE did not interfere with the content learned by students, only in the way it is presented, due to the particularities of each network and the generational difference.

Conducting and directing studies through a single platform, as is still common in the educational environment, is no longer sufficient to encompass the multiplicity of communication currently established^30^, which raises, once again, the importance of verifying whether “Adopt a Bacterium” remains effective on multiple platforms. With the data presented in this work, it’s possible to affirm that the change in social media platforms did not adversely affect student performance, demonstrating that it can be applied across multiple platforms, depending on the teacher’s objectives and the interests and needs of the class. This is crucial when considering social media, as they are constantly being updated and modified, and students change their interests in them at an increasingly rapid pace, requiring planning and adaptation for the project to remain current and thus more closely aligned with students’ reality and able to engage them more effectively.

Due to the years in which the data was collected, a small observation is necessary in relation to the context of remote teaching due to the pandemic period.

In the specific case of remote teaching, which was the context of the 2020 and 2021 #Adopt projects, the main point to be raised concerns the interaction among students and with teachers, as this aspect is significantly impaired in this model^19,20^. However, in the three editions analyzed, it was not possible to verify this decrease in interaction, both among peers and with mediators, which is a very important point, since interaction is an essential part of the teaching-learning process^31^. It was even maintained during the pandemic period. The structure of #Adopt itself promotes this interaction by requiring that work be done in groups at absolutely every stage, including during exams, which already demands closer contact between students, and mediation is an essential part of it, promoting the interaction between mediators and students as well. Furthermore, the environment in which the project takes place, namely social networks, also greatly encourages this interaction, including the teacher-student relationship^13,32,33^. Even when compared to other virtual environments designed from their conception for educational purposes, social networks are more efficient in encouraging interaction and communication among peers^34^.

Therefore, all these observations, combined with the increase in the richness of discourse and the topics mentioned by the students, are strong indications that “Adopt a Bacterium” has proven to be suitable for teaching Microbiology, specifically Bacteriology, even during a period of social isolation and remote learning. The students remained interested and dedicated to their activities, even remotely, which aligns with the general trends observed in education during this period. This only reinforces the importance and potential of “Adopt a Bacterium” as an active methodology for teaching Microbiology.

### The final considerations of this study are as follows

Over the three years of analysis, the #Adopt Project has proven to be effective for student learning, which can be inferred from the richness of discourse being greater in Q2 and Q3 than in Q1 (preconceptions); an increase in the themes mentioned by students in their responses and the frequency of citations of each theme; and a small number of conceptual errors over the three years. Additionally, there was an acquisition of vocabulary in Microbiology. Furthermore, these data demonstrate that #Adopt was able to mitigate many of the effects observed in the pandemic on the education sector, as the data for the three years follow a similar pattern. Moreover, it is capable of sparking students’ interest and motivating them, as demonstrated by the grades and comments made by them about the project. Finally, there is great potential for application across multiple platforms, as there was no significant difference observed between the data obtained in 2020, when Facebook® was used, and the subsequent years analyzed here, when Instagram® became the platform.

## Acknowledgments

This study was financed by the Conselho Nacional de Desenvolvimento Científico e Tecnológico (CNPq), by Coordenação de Aperfeiçoamento de Pessoal de Nível Superior - Brasil (CAPES) - Finance Code 001 and by the “Fundação de Amparo à Pesquisa do Estado de São Paulo (FAPESP) in the CEPID B3 program. We thank all the undergraduate students, graduate students and researchers of the Institute of Biomedical Sciences at the University of São Paulo for collaborating as mediators and elaborating the activities. We also thank the undergraduate students for participating in the project.

